# Tetraploidy accelerates adaption under drug-selection in a fungal pathogen

**DOI:** 10.1101/2021.02.28.433243

**Authors:** Ognenka Avramovska, Emily Rego, Meleah A Hickman

## Abstract

Baseline ploidy significantly impacts evolutionary trajectories, and in particular, tetraploidy has been associated with higher rates of adaptation compared to haploidy and diploidy. While the majority of experimental evolution studies investigating ploidy use *Saccharomyces cerivisiae*, the fungal pathogen *Candida albicans* is a powerful system to investigate ploidy dynamics, particularly in the context of antifungal drug resistance. *C. albicans* laboratory and clinical strains are predominantly diploid, but have also been isolated as haploid and polyploid. Here, we evolved diploid and tetraploid *C. albicans* for ∼60 days in the antifungal drug caspofungin. Tetraploid-evolved lines adapted faster than diploid-evolved lines and reached higher levels of caspofungin resistance. While diploid-evolved lines generally maintained their initial genome size, tetraploid-evolved lines rapidly underwent genome-size reductions and did so prior to caspofungin adaption. Furthermore, fitness costs in the absence of drug selection were significantly less in tetraploid-evolved lines compared to the diploid-evolved lines. Taken together, this work supports a model of adaptation in which the tetraploid state is transient but its ability to rapidly transition ploidy states improves adaptative outcomes and may drive drug resistance in fungal pathogens.

## Introduction

Variation in ploidy (i.e. the number of chromosome sets in a genome) occurs widely across the tree of life with differences between species, within species, and even in different cell types in a single organism (Otto 2007; Otto and Gerstein 2008; Peer et al. 2017). Organisms exists on a ploidy spectrum, from haploidy to polyploidy – with each state offering unique evolutionary benefits and drawbacks based on ploidy specific mutation rates and effect sizes (Mable and Otto 2001; Otto and Gerstein 2008; Gerstein 2013; Selmecki et al. 2015; Sharp et al. 2018; Gerstein and Sharp 2021). Interestingly, compared to haploids and diploids, polyploids generally exhibit much higher levels of genome instability and thus an increased ability to generate mutations (Mayer and Aguilera 1990; Storchová et al. 2006; Hickman et al. 2015; Avramovska and Hickman 2019). A high mutation rate can be advantageous if the limiting factor to adaptation is generating mutations. However, if beneficial mutations are recessive, they will take longest to be unmasked in polyploids (Otto 2007; Peer et al. 2017).

The mechanisms by which ploidy transitions occur varies widely. Organisms with sexual cycles can undergo mating and meiosis to increase and reduce ploidy, respectively. However, asexual organisms can use variations in mitosis or leverage errors during cell division to mediate ploidy transitions. Increases in ploidy through endoreplication, in which DNA duplication occurs but cell division does not, is occurs during development in *Drosophila* larvae and mouse trophoblasts (Peer et al. 2017). has also been linked to increased adaptive potential, such as in human hepatocytes that may be experiencing DNA-damage (Duncan 2013) or in *Arabidopsis thaliana*, in which polyploid root tips led to increased tolerance of salinity compared to diploid tips (Chao et al. 2013). In clonal systems, tetraploidy is a transient state, which occurs through cell-cell fusion events and can be found in cancer cell types with high genomic instability (Matsumoto et al. 2021), in addition to fungal species such as *Cryptococcus neoformans* (Okagaki and Nielsen 2012; Gerstein et al. 2015) and *Candida albicans* (Magee and Magee 2000; Bennett and Johnson 2003; Forche et al. 2008).

In the asexual organisms *Candida albicans*, a baseline diploid, tetraploidy is generated through cell-fusion of cells from opposite mating types (Magee and Magee 2000; Bennett and Johnson 2003; Forche et al. 2008). The tetraploid state is unstable (Hickman et al. 2015) and undergoes random concerted chromosome loss to generate diverse progeny which can carry aneuploidy (Hickman et al. 2015; Hirakawa et al. 2017). In *C. albicans*, aneuploidy and thus copy number variation of resistance-associates genes is a mechanism by which resistance to a commonly used antifungal, fluconazole, arises (Selmecki et al. 2006). Therefore, the ability to undergo ploidy reduction through concerted chromosome loss may be an important mechanism driving adaption in under antifungal-drug selective pressure in asexual eukaryotic organisms.

Due to its non-meiotic genome dynamics and clinical relevance, we used *Candida albicans* to test the hypothesis that tetraploids will adapt faster than diploid in the antifungal drug caspofungin. We chose Caspofungin as the antifungal selective pressure because it is highly mutagenic and fungicidal to both diploid and tetraploid *C. albicans*. To address our hypothesis, we evolved 72 diploid and 72 tetraploid lines in caspofungin for 60 days and tracked adaptation and ploidy throughout evolution. We found that tetraploid-evolved lines adapt faster and reach higher resistance levels than diploid-evolved lines. Furthermore, large scale ploidy reductions occurred prior to adaption, though only in tetraploid-evolved lines. We also measured growth rates in the absence of selection and found ploidy specific differences, with tetraploid incurring half the fitness costs of diploids. Interestingly, we did not detect differences between the growth rates of caspofungin-resistant and caspofungin-resistant populations, suggesting there was no fitness cost to caspofungin resistance. In conclusion, we demonstrate that transient tetraploidy can facilitate adaptation in asexual organisms.

## Materials and Methods

### Yeast strains and media

Stains used in this study are listed in Table S1. All evolved populations were stored in 96-well block format in −80°C and maintained on YPD (1% yeast extract, 2% bactopeptone, 2% glucose, 1.5% agar, 0.004% adenine, 0.008% uridine) or casitone (0.9% bacto-casitone, 0.5% yeast extract, 1% sodium citrate, 2% glucose, 1% agar) media at 30°C. Caspofungin stocks of 1 mg/ml (Sigma-Aldrich CAS#179463-17-3) were made from powder and suspended into ddH_2_0. Liquid yeast cultures were grown in casitone (0.9% bacto-casitone, 0.5% yeast extract, 1% sodium citrate, 2% glucose).

### Experimental Evolution

Diploid strain MH84 and tetraploid strain MH128 were initially struck onto YPD agar plates to obtain single colonies. Experimental evolution was set up with two, 96-well blocks containing YPD, and inoculated with 36 single colonies of MH84 and 36 single colonies of MH128 for each plate and cultured for 24 hrs. The following day, cultures were normalized to 0.5 OD, and 100 uL added to 900uL of casitone supplemented with 0.25 µg/mL caspofungin and 100 mg/mL streptomycin (to prevent bacterial growth) in a 96-well block. Culture blocks were covered with BreathEasy tape and placed in tupperware containers containing damp paper towels to maintain humidity and incubated at 30°C. Every 7 days, 100uL was removed for archival glycerol stocks. The remaining cultures were pelleted by centrifugation (2 min at 1500 RPM), media removed and pellets were resuspended in 1 mL fresh casitone media supplemented with 0.25µg/mL caspofungin and 100 mg/mL streptomycin.

### Relative Caspofungin Growth

Relative caspofungin growth (RCG) was determined every 7 days by spotting 5 uL onto casitone (no-drug) and casitone containing 0.25ug/mL caspofungin (+drug) agar plates. Plates were incubated at 30°C for 24 hrs and subsequently photographed. Photographs were analyzed to determine the pixel cell area for each spot, using Colonyzer imaging software (Lawless et al. 2010). Relative caspofungin growth was calculated by dividing the pixel area of each spot +drug plates by the pixel area into the corresponding no-drug plate and ratios were capped at 1.0, indicating no growth difference in the presence or absence of caspofungin.

### Drug susceptibility

E-test: Minimum inhibitory concentrations (MIC) were measured as previous described (Avramovska and Hickman 2019). Briefly, 10 µL of glycerol stock was inoculated into 2 mL YPD supplemented with 100mg/mL ampicillin and grown with shaking at 30°C for 24hrs. Cultures were normalized to 0.1 OD with ddH_2_0. 200uL was spread onto casitone agar (1%) plates and left to dry for 15 minutes at 30°C. Standardized caspofungin E-test strips (gradient 0.002 µg/mL – 32 µg/mL; Biomeureix) were added to the middle of the plates and incubated at 30° C for 24 hrs and subsequently photographed.

Microbroth Dilution Assay: Microbroth dilution assays were performed as in the CLSI M27-A guidelines with the following modifications. Evolved populations were inoculated into casitone media supplemented with 100mg/mL ampicillin and incubated at 30 °C with shaking for 48 hrs. Cultures were normalized to 1 OD and diluted to yield approximately 1×10^3^ cells/mL. 100uL of the diluted cells were added to 100uL of casitone containing a gradient of caspofungin concentrations (no-drug, 0.03125 ug/mL caspofungin - 4 ug/mL). Blocks were covered with BreatheEasy tape and incubated at 30° C for for 24 hrs. OD600 was measured on a plate reader (BioTekGen5). The drug concentration in which the fraction of growth (relative to no-drug) was below 0.5 was considered the minimum inhibitory concentration.

### Growth rates

Growth rates were determined similar to (Hickman et al. 2015) with some modification. Populations were inoculated from 10 ul of glycerol stock into 490uL of YPD with 100 mg/mL ampicillin and grown for 24hrs with shaking at 30C to ensure that cells have recovered. Following 24 hr growth, 1:200 dilution was performed into fresh YPD media and OD600 was measured every 15 minutes with shaking using BioTek5 growth reader. Growth rate were determined using R-script (Gerstein et al. 2012).

### Flow cytometry

Flow cytometry analysis was performed as previously published (Hickman et al. 2015; Avramovska and Hickman 2019). Initially, 200uL of cells in midlog-phase were harvested, washed with distilled water, and resuspended in 20uL of 50 mM Tris (pH 8):50 mM EDTA (50:50 TE). Cells were fixed with 95% ethanol and incubated at 4°C overnight. Following ethanol fixation, cells were washed twice with 50:50 TE, resuspended in 50µL of 1 mg/ml RNAse A and incubated at 37°C for 1-3hrs. Cells were then collected, resuspended in 50µL of 5 mg/ml Proteinase K, and incubated at 37°C for 30mins. Cells were subsequently washed once with 50:50 TE, resuspended in 50µL SybrGreen (1:100 of dilution in 50:50 TE), (Lonza, CAT#12001-798, 10,000x concentrated) and incubated overnight at room temperature. Cells were then collected via centrifugation and resuspended in 150 µL 50:50 TE, briefly sonicated, and run on a LSRII machine with laboratory diploid (MH1) and tetraploid (MH2) strains serving as calibration and internal controls. To estimate the average G1 peak FITC-A or BB515 intensity, the multi-Gaussian cell cycle model was used (FloJoV10).

### Statistical analysis

Statistical analysis was performed using GraphPad Prism 9 software.

## Results

### Tetraploids evolve faster and attain higher antifungal resistance than diploids

There are only a limited number of experimental evolution studies that have investigated how tetraploidy impacts the rate and magnitude of adaptation (Gerstein et al. 2006, 2008, 2017; Selmecki et al. 2015) and even fewer that have investigated how tetraploidy contributes to the emergence of antifungal drug-resistance in fungal pathogens (Okagaki et al. 2010; Harrison et al. 2014). We and others have previously shown that tetraploids have both higher baseline and drug-induced genome instability compared to diploids (Hickman et al. 2015; Avramovska and Hickman 2019), thus we hypothesize that tetraploidy facilitates rapid adaptation to antifungal drugs. To test this hypothesis, we compared 72-diploid and 72-tetraploid replicate *C. albicans* lines evolved in the antifungal drug, caspofungin. We evolved the replicate lines in 0.25 µg/mL caspofungin, a concentration that reduces viability by 99% (Avramovska and Hickman 2019), for 59 days. Given the fungicidal nature of caspofungin, replicate lines were not subjected to bottlenecks, instead selective media was replenished every week. Throughout the course of experimental evolution, we simultaneously spotted replicate lines onto selective and non-selective media and photographed 24 hrs later for image analysis (Fig. 1A). For each replicate line, we calculated the relative caspofungin growth (RCG) by dividing the area of growth on selective media by the area of growth on non-selective media for days 0, 3, 10, 18, 27, 37, 45, 52 and 59 (Fig 1A, B and S1). Detectable RCG changes were initially observed on day 18, with average RCGs of 0.02 and 0.21 for diploid- and tetraploid-evolved lines, respectively. Day 18 tetraploid-evolved RCG was largely driven by 19 replicate lines greater than 0.5 (Fig S1B). In contrast, only a single diploid-evolved line had an RCG larger than 0.5 at this timepoint. Throughout the evolution, tetraploid-evolved lines had significantly higher average RCG than diploid-evolved lines (Fig.1B). By day 59, tetraploid-evolved RCG was 0.68 compared to the 0.33 diploid-evolved RCG. This data supports the hypothesis that tetraploids not only adapt more rapidly than diploids, they improve by significantly larger margins.

**Figure 1.**
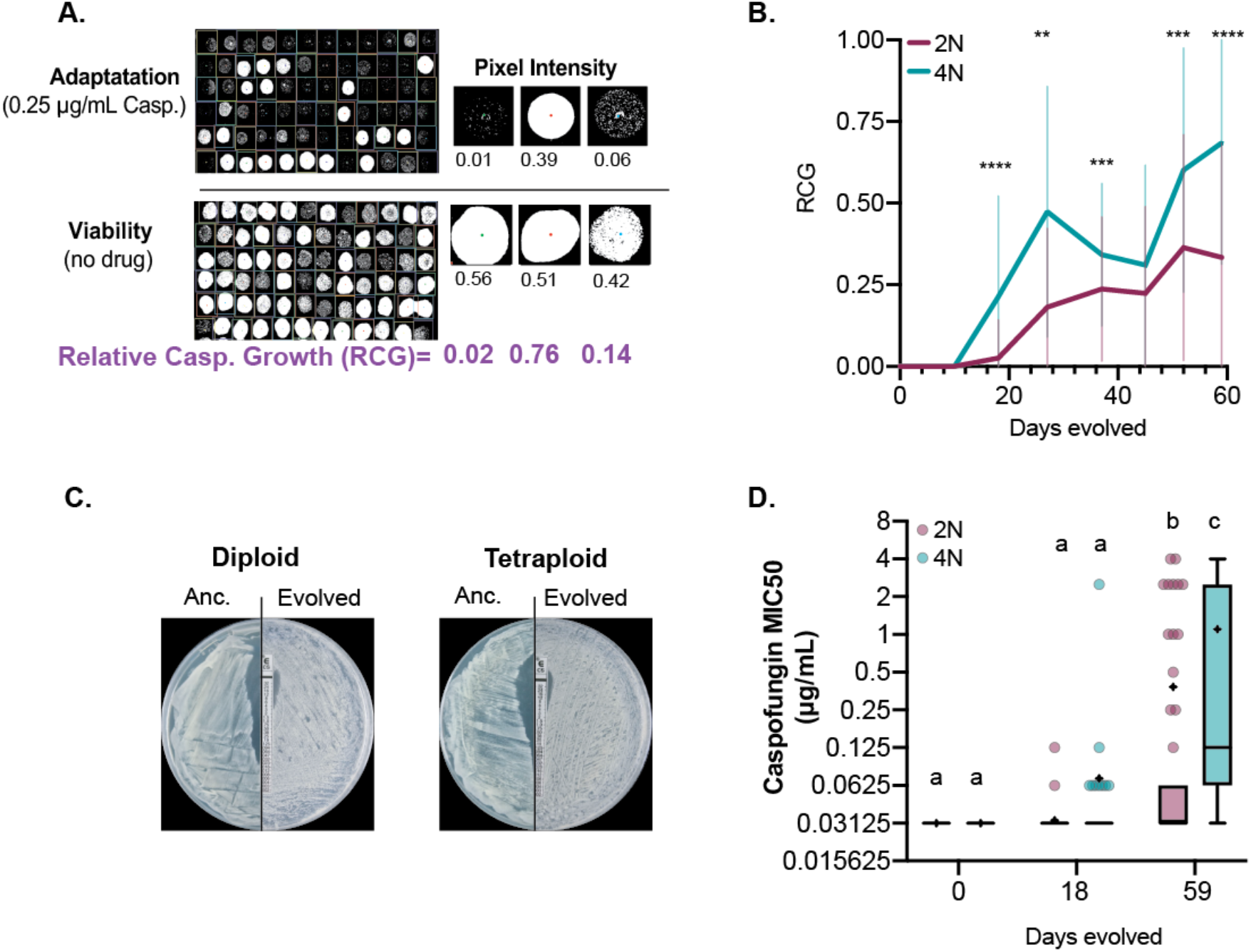
Diploid and tetraploid *Candida albicans* evolution under caspofungin selection. **A)** Experimental setup of 72 diploid- and 72-tetraploid evolved lines exposed to 0.25 μg/mL for 59 days. Relative caspofungin growth (RCG) was determined by spotting evolved lines on no-drug and caspofungin (0.25μg/mL) plates, photographed after 24 hrs and pixel intensity measured for each spot to determine the ratio of caspofungin growth to total viable growth. Images are of well F8, F9 and F10 imaged on Day45 of experimental evolution. Significant differences between diploid and tetraploid evolved lines per timepoint were tested using Mann-Whitney U-test, **** p<0.0001, *** p<0.0001, ** p<0.005. **B)** Average RCG for diploid (maroon; n=72) and tetraploid (teal; n=72) replicate lines over 59 days of caspofungin evolution. RCG was measured on Days 0, 3,10,18,27,37,45,52,59. Error bars represent +/− 1SD. **C)** Representative e-tests of ancestral and evolved (Day 59) diploid (line 64) and tetraploid (line 35). **D)** Caspofungin minimum inhibatory concentration (MIC50μg/mL) of diploid- and tetraploid-evolved lines for Day0, Day18, and Day59. Data is displayed as Tukey-box-and-whisker plots, the mean is indicated by (+) and median is dark black line and circles indicate outliers. Statistical analysis is Kruskal-Wallis test with Dunn’s post hoc multiple-comparison testing. Statistical significance is designated by letters that differ from one another.

While RCG captures caspofungin adaptation over the course of experimental evolution, it may not necessarily reflect caspofungin resistance, a clinical term defined as a minimum inhibitory concentration (MIC) of 1µg/mL (Santos et al. 2014). MIC is measured either by growth on agar plates with a drug concentration gradient (i.e. e-tests, Fig. 1) or in liquid culture as a microbroth dilution assay. We measured MIC on days 0, 18, and 59 for all replicate lines (Fig. 1D). There were significant differences in MIC between diploid- and tetraploid-evolved lines, despite ancestral MIC being ∼0.03 for both ploidy states. By day 18, the diploid-evolved MIC had not changed from the ancestral, whereas tetraploid-evolved MIC had increased to 0.07 ug/ml. By day 59, tetraploid-evolved MIC was still significantly higher (1.0µg/ml) than diploid-evolved MIC (0.4 ug/ml), a pattern consistent with RCG values (Kruskal-Wallis, p<0.0001, Table S2).

We assessed the relationship between MIC and RCG, and found that evolved-lines with MIC values ≥ 0.25 µg/ml caspofungin have RCG values 0.71 or greater. However, this relationship breaks down for evolved lines with MICs less than 0.25 µg/ml (Fig. S2). Therefore, we measured changes in MIC, rather than RCG in subsequent analyses. By day 18, 13% (9/72) of tetraploid-evolved lines, and 4% (2/70) diploid-evolved lines had increased MIC by at least two-fold (Fig. S2A & B). By the end of the evolution experiment, nearly all the tetraploid-evolved lines (70/72) had increased MIC, whereas only half of the diploid-evolved lines (35/72) had improved (Fig. S2C & D). Interestingly, 10% (7/72) of diploid-evolved lines and nearly 40% (28/72) of tetraploid-evolved lines had MIC values with a ten-fold or higher increase in drug concentration than the selective pressure they were evolved in (Fig. S2C & D). Regardless of how growth in caspofungin was quantified, we found that tetraploids evolved faster than diploids and more frequently attained drug-resistant phenotypes.

### Large scale genome size reductions occur prior to adaptation

*C. albicans* tetraploid genomes are intrinsically unstable and frequently undergo chromosome loss to return to a diploid, or near-diploid state over time (Bennett and Johnson 2003; Forche et al. 2008; Hickman et al. 2015; Gerstein et al. 2017). We have previously shown that short-term exposure to caspofungin rapidly induces chromosome loss in tetraploids(Avramovska and Hickman 2019) and thus, we next investigated whether there was any relationship between changes in genome size and adaptation to caspofungin. Throughout the experimental evolution, we measured the genome size for all diploid- and tetraploid-evolved lines (Fig. 2A&B and Fig. S3, Table S3) and observed that tetraploids reduced extensively in genome size within the first 10 days in contrast to diploids, which showed slight, but statistically significant deviations. By day18, diploid-evolved and tetraploid-evolves lines have approximately the same genome-size, suggesting that tetraploid-evolved lines have reached a near diploid state (Fig. 2A&B). Next, we plotted the changes in genome size by the changes in MIC for diploid- and tetraploid-evolved lines for day 18 (Fig 2C) and for day 59 (Fig 2D). By day18 diploid-evolved lines displayed either no change or modest increases in genome size, and only 2 lines had increased MIC (Fig 2C). By day 59, 35 diploid-evolved lines increased in MIC, but had not changed in genome size. In contrast, by day18, most tetraploid-evolved lines reduced in genome size, with a majority near diploid, yet only nine lines had increased MIC. By day 59, nearly all tetraploid lines had increased MIC, but had not undergone additional genome size changes after day18 (Fig 2D). From this data, we conclude that while tetraploids rapidly reduce in genome size, this occurs prior to caspofungin adaptation and there is no clear relationship between genome size and caspofungin resistance phenotypes.

**Figure 2:**
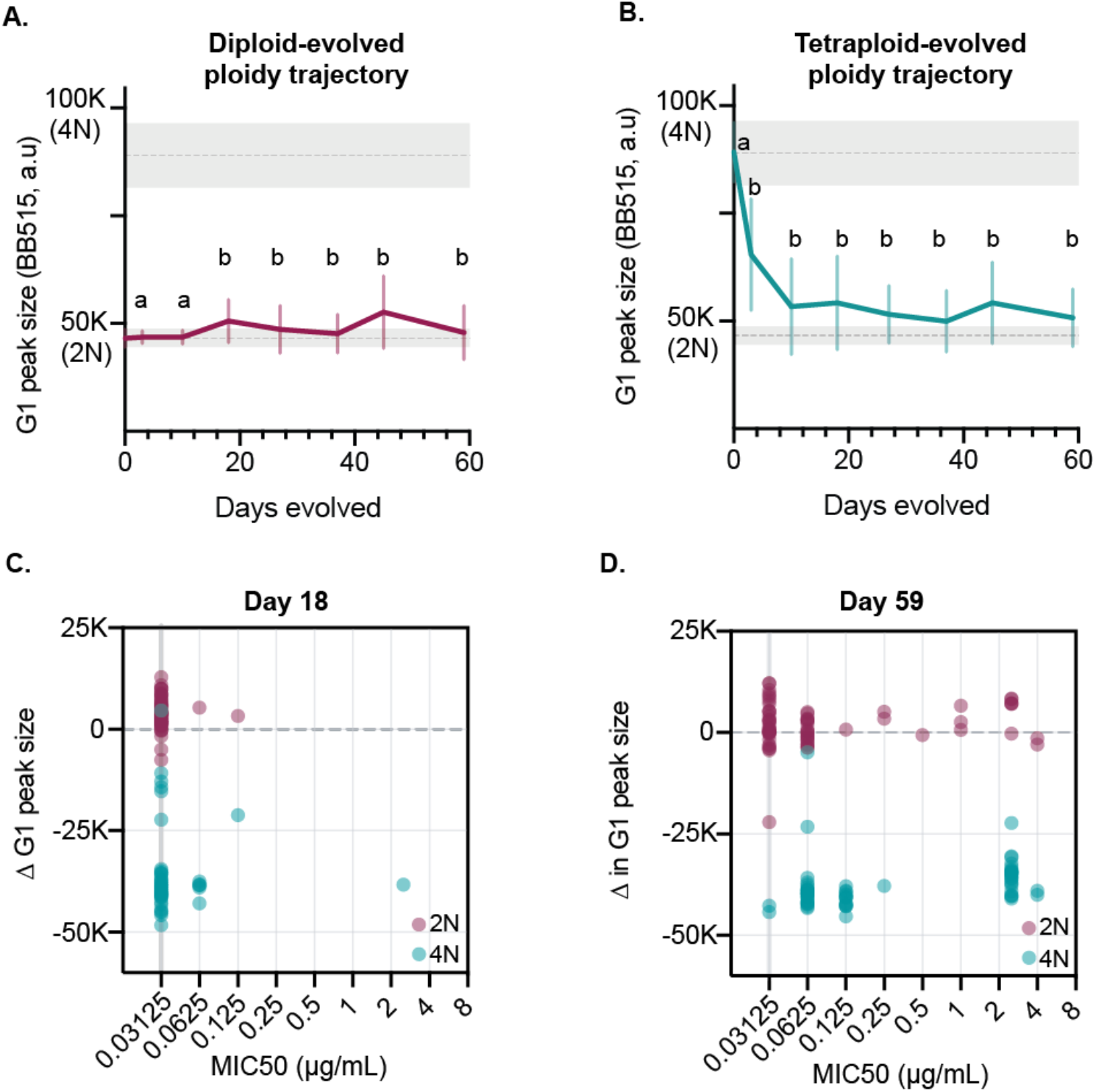
Large scale genome size reductions occur prior adaption. **A)** Diploid genome size measured throughout experimental evolution (day 3,10,18,27,37,45,59) using flow cytometry with the mean and SD of G1 peak plotted in a.u (arbitrary units). The gray lines represent the mean of the ancestral diploid control (n = 72). Each timepoint represents the mean genome size of at least 10,000 events per replicate line (n) and includes G1 peaks of mixed ploidy population. Statistical comparisons are between evolved and ancestral (D0) and statistical significant is determined by Kruskal-Wallis with posthoc Dunn’s multiple comparison. Timepoints designated with ‘b’ are significantly different from ancestral. **B)** Tetraploid genome size measured throughout experimental evolution. Method and statistical comparison is same as in ‘A’. **C)** Change in genome size by Day 18 compared to ancestral diploid and tetraploid averages is plotted on y-axis. MIC50 is plotted on the x-axis for each replicate line, with the acestral MIC designated by the gray vertical line at 0.03125μg/mL. **D)** Change in genome size by Day 59 compared to ancestral diploid and tetraploid averages is plotted on y-axis. MIC50 is plotted on the x-axis for each replicate line, with the acestral MIC designated by the gray vertical line at 0.03125μg/mL.

### Tetraploids have lower fitness costs associated with evolution under caspofungin selection

During the caspofungin experimental evolution, we found that growth in the absence of drug selection was highly variable, with some lines growing robustly and others barely viable (Fig. 1A, no-drug), likely from mutation accumulation induced by caspofungin (Avramovska and Hickman 2019). To test whether caspofungin evolution results in reduced fitness in the absence of drug (no-drug), we measured the growth rate for ancestral (Day0) and evolved (Day59) replicate lines in the absence of caspofungin (Fig. S4A). While there was a fitness cost for both diploid- and tetraploid-evolved lines, the diploid cost (ΔGR = −0.08) was twice as large as the tetraploid (ΔGR= −0.04) (Fig. 3A). However, the initial growth rates of tetraploid lines was lower than diploid lines (p < 0.0001 Mann-Whitney U-test, Fig. S4A) and there was a significant negative relationship between initial and evolved growth rates (Fig. S4B, R^2^ = 0.41, p < 0.0001); lines with higher initial growth rates had greater reductions in growth rates following caspofungin evolution.

**Figure 3.**
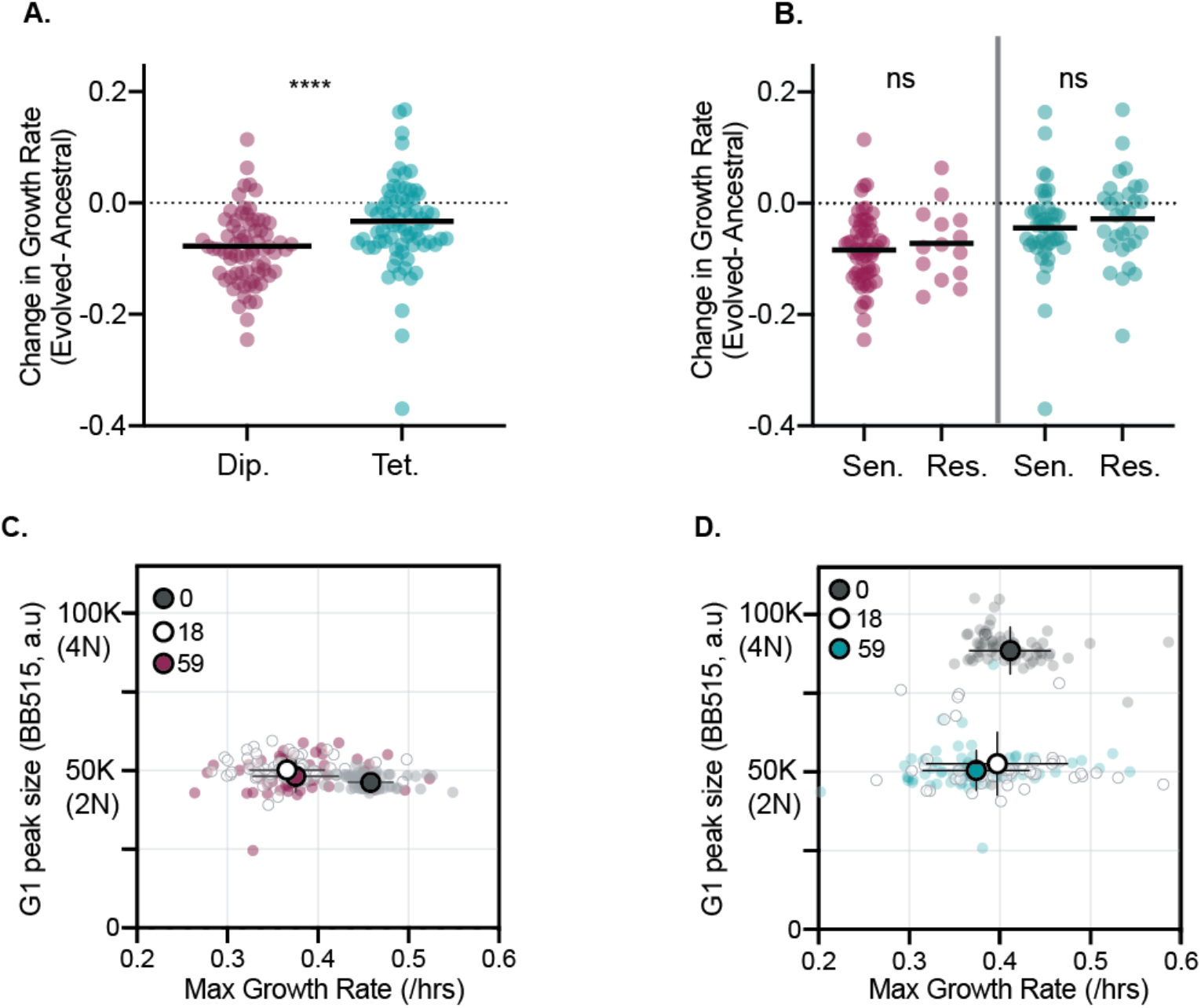
Tetraploids have lower fitness costs associated with evolution under caspofungin selection. **A)** The change in growth rate (D59-D0) for each diploid (maroon) and tetraploid (teal) evolved line is plotted, with the dark black line respresenting the mean. Statistical comparison between diploid and tetraploid is represented by Mann-Whitney U-test (****, p<0.0001). **B)** The change in growth rate (D59-D0) for each sensitve and resistant diploid (maroon) and tetraploid (teal) evolved line is plotted, with the dark black line respresenting the mean. Statistical comparison is between sensitive and resistant populations of the same ploidy state and is represented by Mann-Whitney U-test. **C)** Growth rate (x-axis) and genome size (y-axis) were plotted for all diploid-evolved replicate lines on day 0 (gray), day 18 (white) and day 59 (maroon). The circles outlined in black represent the mean for each time point and black vertical and horizontal lines represent standard deviation. **D)** Growth rate (x-axis) and genome size (y-axis) were plotted for all tetraploid-evolves replicate lines on day 0 (gray), day 18 (white) and day 59 (teal). The circles outlined in black represent the mean for each time point and black vertical and horizontal lines represent standard deviation.

To test if the reductions in no-drug growth rates were a trade-off with caspofungin resistance, we compared growth rates of evolved lines with an MIC greater than or equal to 1.0 µg/ml caspofungin (‘resistant’), the clinical concentration for a strain to be considered drug-resistant, to the evolved lines with an MIC less than 1µg/ml (‘susceptible’) in the absence of drug (Fig. 3B). For both diploid- and tetraploid-evolved lines, there was no difference in no-drug growth rates between resistant and susceptible evolved lines (Mann-Whitney U-test p= 0.99). Thus, reductions in no-drug growth rates were not due to a direct trade-off with resistance to caspofungin. Rather, reductions in no-drug growth rate are likely due to the accumulation of deleterious caspofungin-induced mutations, including genome-size changes. We next measured no-drug growth rates on day18, a timepoint with minimal detectable caspofungin adapatation, yet significant genomes-size changes (Fig 3C & D). In diploid-evolved lines, by day18 there was significant reduction in no-drug growth rates compared to day 0 (p <0.0001, Friedman test, Table S4), but no further reduction in growth rates by day 59 (p>0.99, Friedman test, Table S4). However, in tetraploid-evolved lines, day18 growth rates were comparable to day 0 (p>0.99, Friedman test, Table S4) despite massive reductions in genome size. By day 59, no-drug growth rates were slower than day 18 (p= 0.037, Friedman test), yet no additional genome size changes occurred. From these results, we propose that tetraploidy buffers against the early fitness costs of drug-induced mutagenesis by purging chromosomes with deleterious mutations. However, once tetraploid genomes have reduced to diploid or near-diploid states, they are subject to carrying high mutational loads that results in fitness costs in the absence of selection.

## Discussion

In this study, we evolved diploid and tetraploid *C. albicans* to the antifungal drug caspofungin to compare how ploidy impacts evolutionary trajectories, and found tetraploids adapted more rapidly and achieved higher levels of drug resistance compared to diploids. Early in the experiment (Day18) tetraploid-evolved MIC was twice that of ancestral, yet diploid-evolved lines showed no changes in MIC (Fig. 1). By the end of the evolution tetraploid-evolved lines improved their MIC by 32-fold, compared to diploid-evolved lines that improved only by 2.5-fold. Our work demonstrates that tetraploidy can facilitate adaptation, a result that is consistent with other experimental evolution studies investigating the role of ploidy in adaptive processes. For example, *S. cerevisiae* tetraploids evolved under raffinose selection also adapted at a significantly faster rate than diploids (Selmecki et al. 2015). In part, accelerated adaptation may be due to elevated mutation rates observed in polyploids relative to diploids across yeast species (Mayer and Aguilera 1990; Storchová et al. 2006; Hickman et al. 2015), leading to higher frequencies of beneficial mutations (Selmecki et al. 2015).

In addition to higher mutation rates, tetraploid cells also frequently undergo random chromosome loss and reduce in genome size under a diverse set of growth conditions *in vitro* and *in vivo* (Bennett and Johnson 2003; Forche et al. 2008; Hickman et al. 2015; Gerstein et al. 2017; Avramovska and Hickman 2019; Smith and Hickman 2020). Interestingly, we found that many of the tetraploid-evolved lines rapidly reduced in genome size by day 10, a timepoint prior to when we first observed any detectable caspofungin adaptation (Figure 2A). Despite returning to diploid (or near-diploid) genome sizes, tetraploid-evolved lines still adapted more quickly and with larger increases in MIC than diploid-evolved lines. Non-meiotic ploidy reduction in *C. albicans* increases phenotypic variation amongst isolates derived from tetraploid cells (Hickman et al. 2015; Hirakawa et al. 2017). We propose that tetraploids are capable of accelerated adaptation because ploidy reduction generates derivatives whose chromosomes have been reassorted, thus carrying new combinations of alleles, and may also contain chromosomal aneuploidy (Forche et al. 2008; Hickman et al. 2015; Hirakawa et al. 2017).

Aneuploidy is considered to be a ‘quick-fix’ that organisms use during adaptation by altering gene expression and protein abundance (Rancati et al. 2008; Pavelka et al. 2010; Yona et al. 2012). Specifically, chromosomal aneuploidy drives heat-tolerance in *Saccharomyces cerevisiae* (Yona et al. 2012), chemotherapy resistance in cancer cells (Duesberg et al. 2000; Gordon et al. 2012) and drug resistance in fungal pathogens (Selmecki et al. 2006, 2009; Sionov et al. 2010; Harrison et al. 2014; Todd et al. 2017; Stone et al. 2019). For example, in *Cryptococcus neoformans*, aneuploidy of chromosomes 1 is associated with resistance to the antifungal drug fluconazole (Sionov et al. 2010) and in *C. albicans* and *C*.*auris*, chromosome 5 aneuploidy confers fluconazole resistance (Selmecki et al. 2006, 2009; Bing et al. 2020). While aneuploidy is an adaptative mechanism for fluconazole resistance in a broad range of fungal pathogens, it has not yet been observed in caspofungin or other echinocandin antifungal drugs. In this study, we found large-scale genome reduction in tetraploid lines and detected smaller-scale increases in genome size in the diploid-evolved lines, indicative of chromosomal aneuploidy across both ploidy states. However, flow cytometry can only measure total genome size and does not have the resolution to identify the karyotypic composition. Further genome characterization via whole-genome sequencing is needed to identify the specific mutations driving caspofungin resistance from this experimental evolution.

For some organisms, acquiring resistance to drugs confers a fitness cost in the absence of drug selection (Dahlberg and Chao 2003; Andersson and Hughes 2010; Vincent et al. 2013; Hill et al. 2015; Popp et al. 2017). In our study, we found no difference in the growth rates between caspofungin susceptible and resistant evolved lines when grown in the absence of drug (Fig. 3B), suggesting that caspofungin resistance may not explicitly confer a fitness tradeoff. However, we found that on average, both diploid- and tetraploid evolved lines had growth deficits compared to their ancestral state, in the absence of caspofungin (Fig. 3A & S3). Surprisingly, the magnitude of the growth deficit depended on the initial ploidy state, with diploid-evolved lines exhibiting twice the deficit of tetraploid-evolved lines. Given the mutagenic nature of caspofungin (Shields et al. 2018; Avramovska and Hickman 2019), it is likely that caspofungin exposure generated high mutational loads during evolution that are expressed when drug selection is removed. However, since tetraploid-evolved lines undergo genome size reductions within the first 10 days of caspofungin evolution, in which chromosomes are stochastically lost (Hickman et al. 2015), deleterious mutations can be quickly removed from the population. In fact, we see no fitness costs for tetraploid-evolved lines at day 18, following genome size reductions (Fig. 3D). In contrast, diploid-evolved lines have no clear mechanisms for purging mutations that are deleterious in the absence of selection, and we see significantly slower no-drug growth rates on day18 compared to the ancestral state (Fig. 3C).

In conclusion, we propose that tetraploidy is a transient state with high adaptative potential compared to diploidy. This work, along with other experimental evolution studies using yeast species show that asexual whole-genome ploidy transitions occur frequently during short- and long-term evolution (Mable and Otto 2001; Gerstein et al. 2006, 2008, 2017; Hickman et al. 2013, 2015; Selmecki et al. 2015; Turanli-Yildiz et al. 2017; Harari et al. 2018). These findings, coupled with the observation of ploidy variability within clinical and environmental isolates of various yeast species (Ford et al. 2015; Hirakawa et al. 2015; Wertheimer et al. 2016; Zhu et al. 2016; Ropars et al. 2018; Stone et al. 2019; Gerstein and Sharp 2021; Scopel et al. 2021) indicate that ploidy transitions may be an important evolutionary force driving microbial eukaryotic adaptation.

## Supplemental

**Figure S1:**
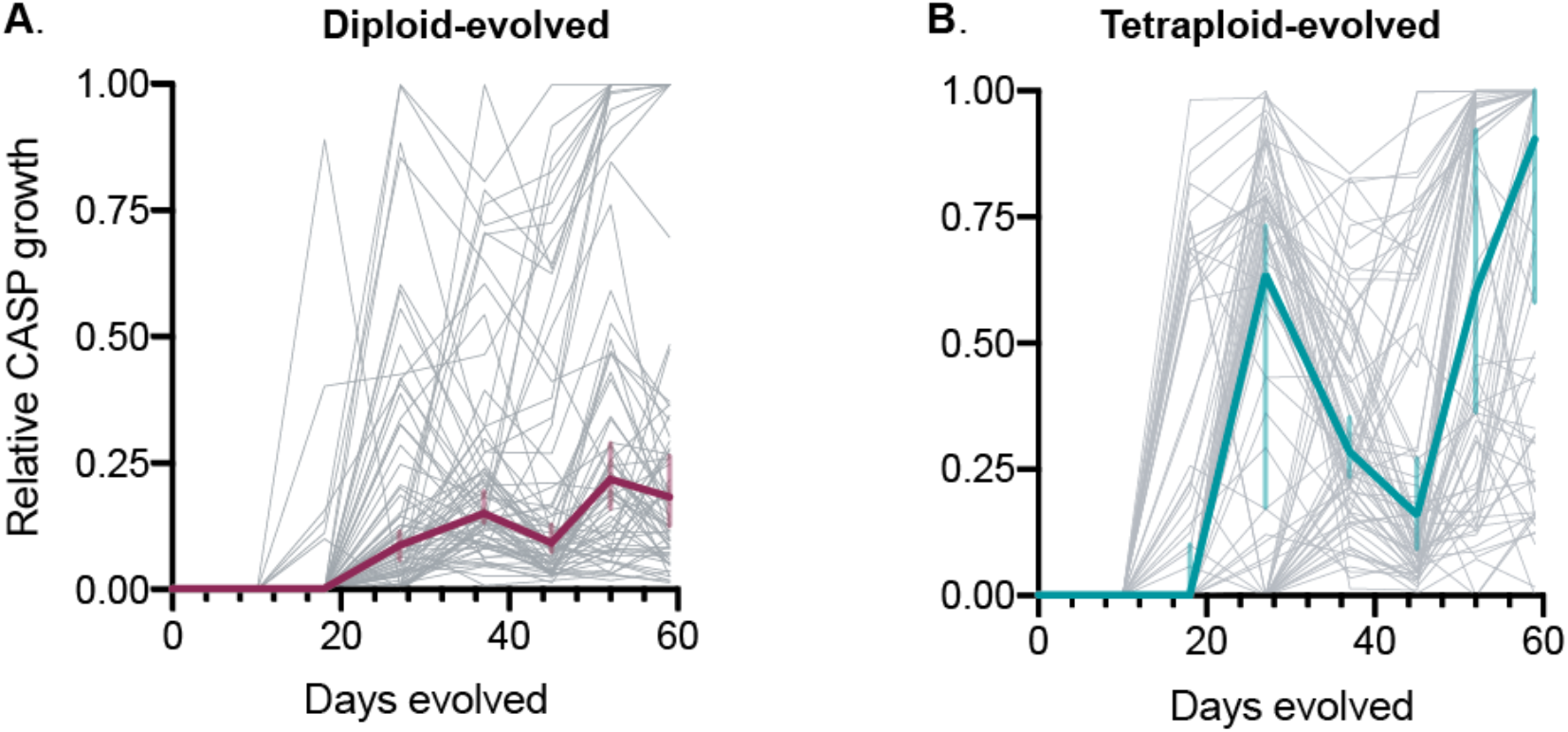
Variation among replicate lines in RCG and MIC values. **A)** Diploid evolutionary trajectories of relative caspofungin growth (RCG) over time. Light gray lines represent trajectories of individual replicate lines (n=72). Maroon line represent the median RCG at the timepoint, with the maroon vertical lines representating the 95% confidence intervals. **B)** Tetraploid evolutionary trajectories of relative casp growth (RCG) over time. Light gray lines represent trajectories of individual replicate lines (n=72). Teal line represent the median RCG at the timepoint, with the teal vertical lines representating the 95% confidence intervals.

**Figure S2:**
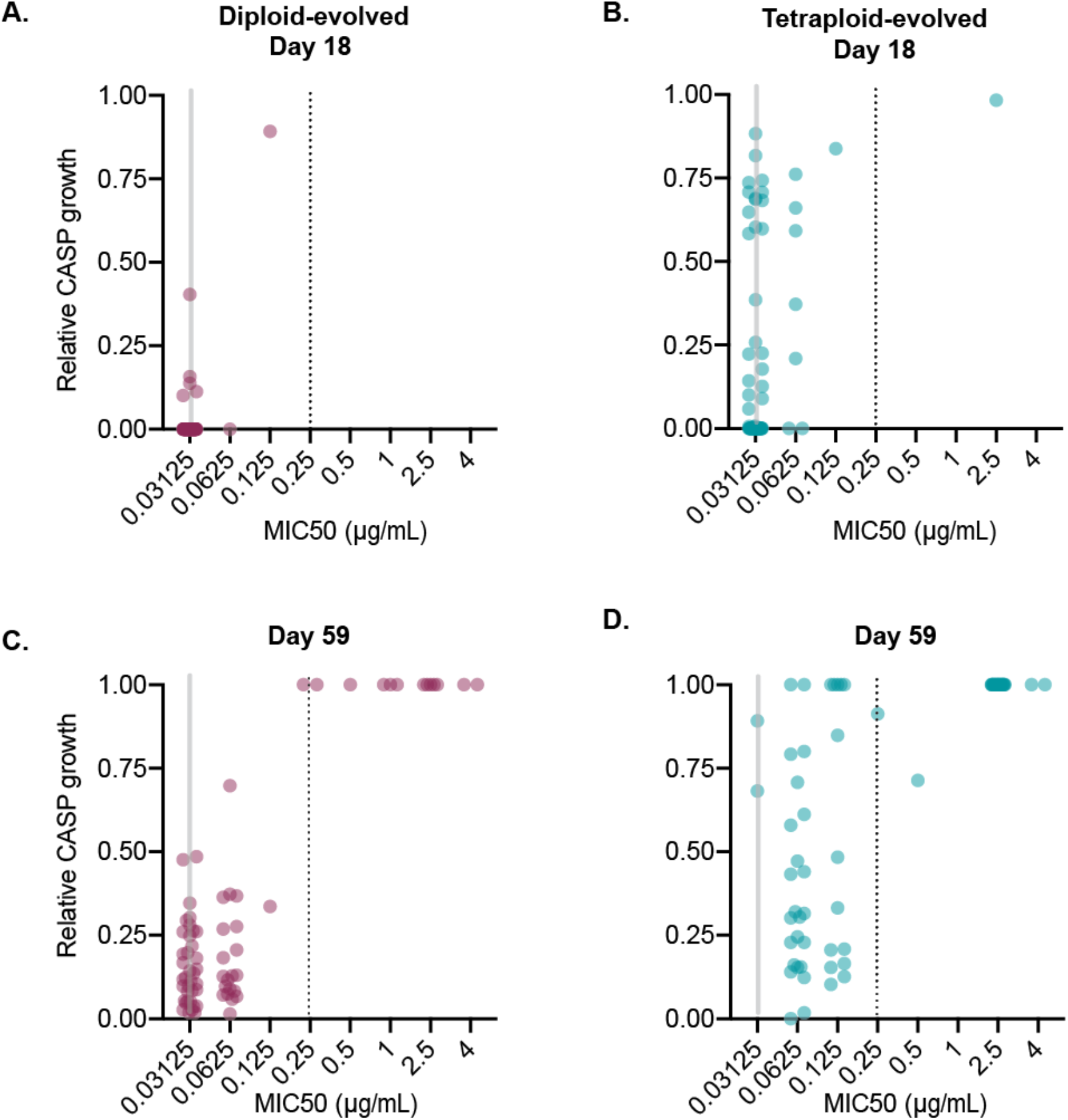
Relative caspifungin growth is associated with MIC, but only at high RCG values. **A)** Relative caspofungin growth and MIC50 values are plotted for diploids (maroon, n=71) and tetraploids (teal, n=70) day 18. Gray line indicates ancestral MIC and black dashed line indicates the concentration of the caspofungin selective pressure 0.25μg/mL. **B)** Relative caspofungin growth and MIC50 values are plotted for diploids (maroon, n=71) and tetraploids (teal, n=70) day 18. Gray line indicates ancestral MIC and black dashed line indicates the concentration of the caspofungin selective pressure 0.25μg/mL. **C)** Relative caspofungin growth and MIC50 values are plotted for diploids (maroon, n=72) and tetraploids (teal, n=72) day 59. Gray line indicates ancestral MIC and black dashed line indicates the concentration of the caspofungin selective pressure 0.25μg/mL. **D)** Relative caspofungin growth and MIC50 values are plotted for diploids (maroon, n=72) and tetraploids (teal, n=72) day 59. Gray line indicates ancestral MIC and black dashed line indicates the concentration of the caspofungin selective pressure 0.25μg/mL.

**Figure S3:**
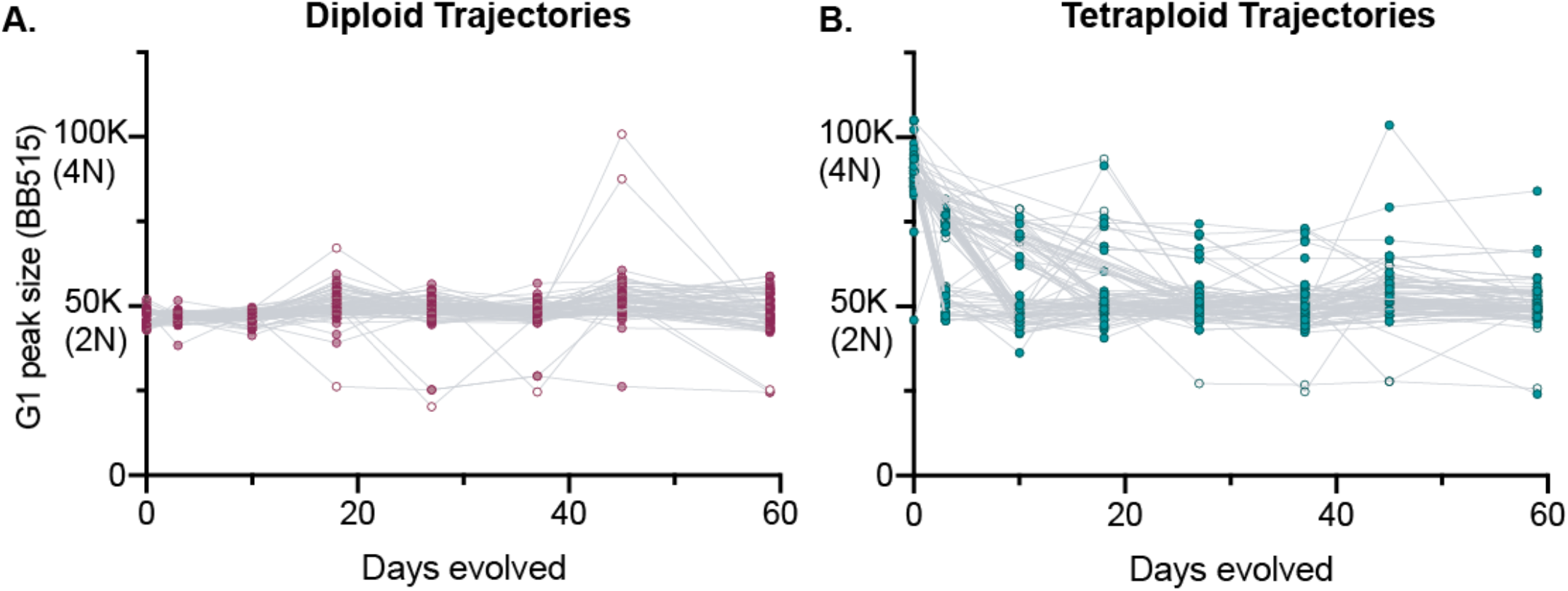
Tetraploid-evolved lines rapidly undergo genome-size reductions during evolution. **A)** Diploid-derived (maroon) genome size measured throughout experimental evolution using flow cytometry (Days 3,10,18,27,37,45,59) with the G1 peak of each replicate line plotted in a.u (arbitrary units). Each symbol represents the mean genome size of at least 10,000 events per population. Open circles represent the G1 peaks of mixed ploidy population (timepoints: Diploid-day3, n = 72; day10, n = 65; day18, n = 65; day27, n = 72, day 37, n = 73; day45, n = 69, day59, n = 71 **B)** Tetraploid-derived (teal)genome size measured throughout experimental evolution using flow cytometry (Days 3,10,18,27,37,45,59) with the G1 peak of each replicate line plotted in a.u (arbitrary units). Each symbol represents the mean genome size of at least 10,000 events per population. Tetraploid - day3, n = 47; day10, n = 57; day18, n = 55; day27, n = 73, day 37, n = 76; day45, n = 57, day59, n = 73).

**Figure S4:**
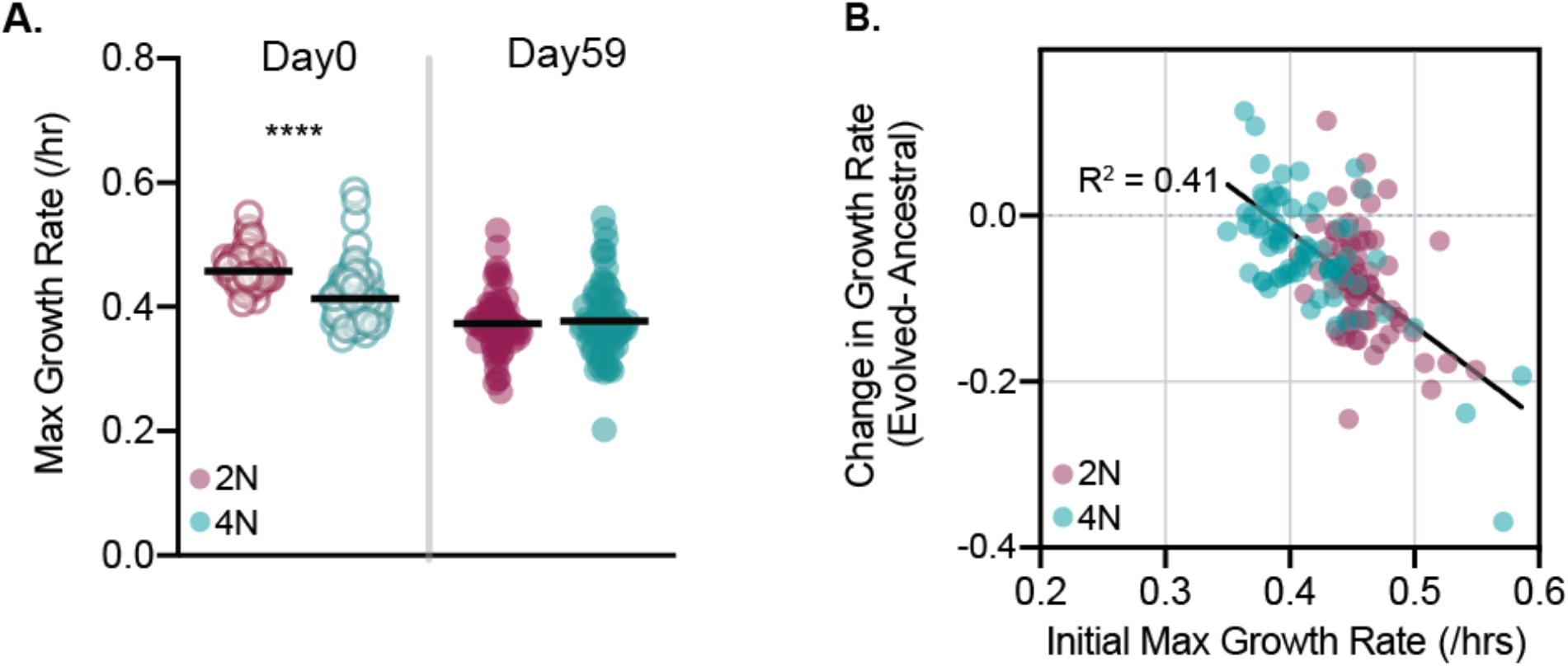
Caspofungin-evolution leads to decreases of growth rates in no-drug conditions. **A)** Maximal growth rate for ancestral diploid and tetraploid lines (D0) and evolved lines (D59) is plotted. Black horizontal lines represent the mean. Statistical testing in comparison between diploid and tetraploid on the same day (Mann-Whitney U-test, ****p<0.0001. **B)** Initial (D0 growth rate is plotted on the x-axis, and the difference between D59 and D0 is plotted on the y-axis for plotted all diploid (maroon, n=72) and tetraploid-evolved lines (teal, n=72). R^2^ value is 0.41, and the slope is statistically significant non-zero (p<0.0001)

**Table S1:**
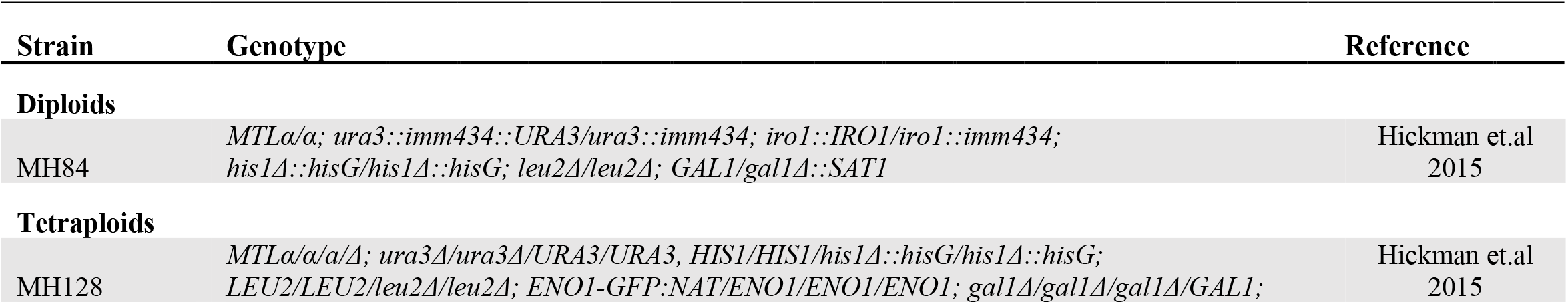
*Candida albicans* strains used in this study

**TS2:**
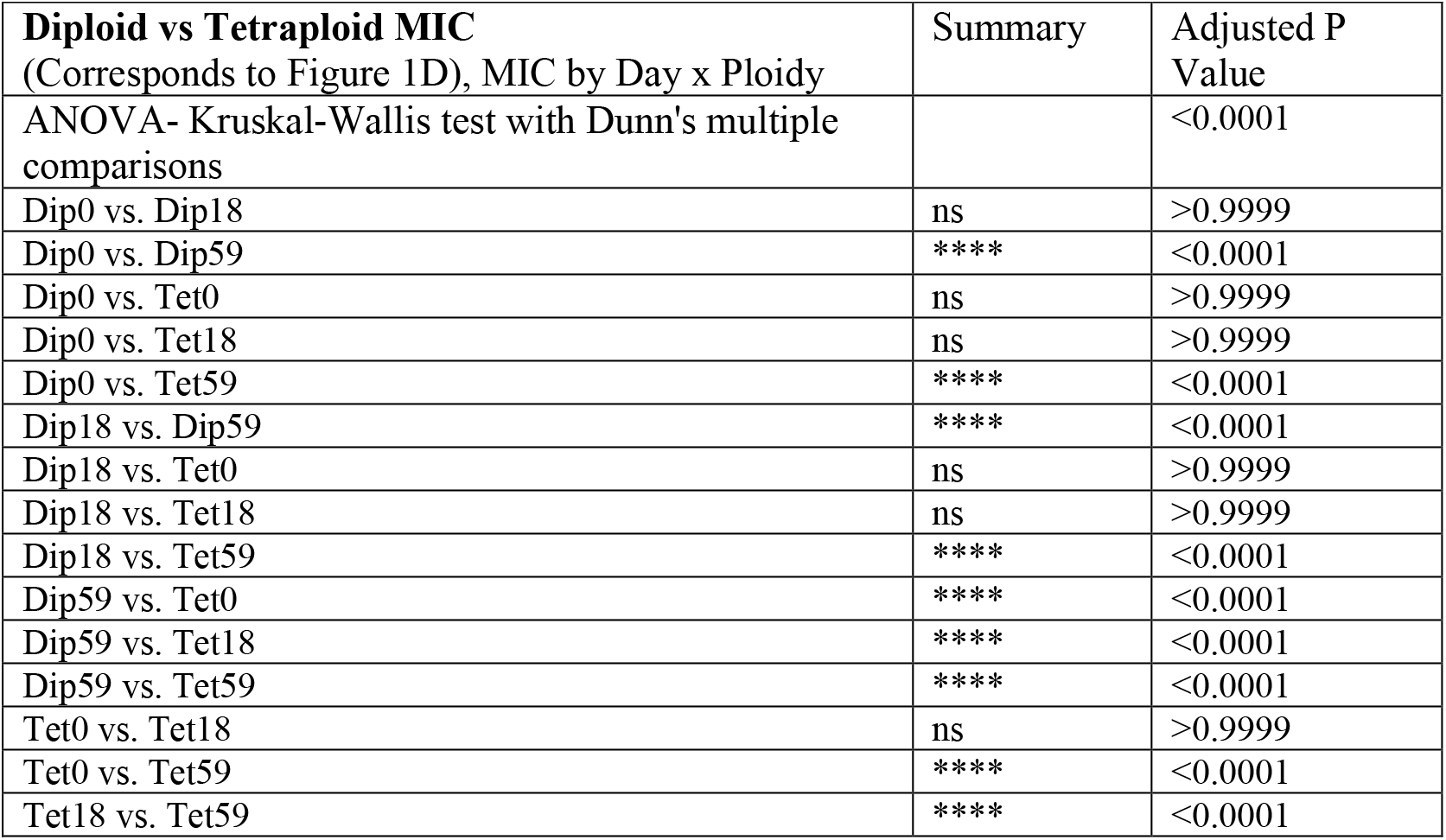
Statistical summaries for figure 1D – multiple comparisons testing

**TS3:** Statistical summaries for figure 2 – multiple comparison’s testing

|  |  |  |
| --- | --- | --- |
| **Diploid Ploidy trajectories**<br>(Corresponds to Figure 2A) | Summary | Adjusted P Value |
| ANOVA- Kruskal-Wallis test with Dunn's multiple comparisons |  | <0.0001 |
| D0 vs. D3 | ns | >0.9999 |
| D0 vs. D10 | ns | >0.9999 |
| D0 vs. D18 | **** | <0.0001 |
| D0 vs. D27 | **** | <0.0001 |
| D0 vs. D37 | ** | 0.0027 |
| D0 vs. D45 | **** | <0.0001 |
| D0 vs. D59 | ** | 0.0064 |
| **Ploidy Ploidy trajectories**<br>(Corresponds to Figure 2B) | Summary | Adjusted P Value |
| ANOVA- Kruskal-Wallis test with Dunn's multiple comparisons |  | <0.0001 |
| D0 vs. D3 | **** | <0.0001 |
| D0 vs. D10 | **** | <0.0001 |
| D0 vs. D18 | **** | <0.0001 |
| D0 vs. D27 | **** | <0.0001 |
| D0 vs. D37 | **** | <0.0001 |
| D0 vs. D45 | **** | <0.0001 |
| D0 vs. D59 | **** | <0.0001 |

**TS3: Statistical summaries for figure 2 – multiple comparison's testing**

**TS4:** Statistical summaries for figure 3 ╌ multiple comparisons testing

|  |  |  |
| --- | --- | --- |
| **Diploid no-drug growth rates** (Correspond to Figure 3C) | Summary | Adjusted P Value |
| ANOVA- Friedman test with Dunn's multiple comparisons |  | p <0.0001 |
| Dip0 vs. Dip18 | **** | <0.0001 |
| Dip0 vs. Dip59 | **** | <0.0001 |
| Dip18 vs. Dip59 | ns | >0.9999 |
| **Tetraploid no-drug growth rates** (Correspond to Figure 3D) | Summary | Adjusted P Value |
| ANOVA- Friedman test with Dunn's multiple comparisons |  | p = 0.0026 |
| Tet0 vs. Tet18 | ns | >0.9999 |
| Tet0 vs. Tet59 | ** | 0.0029 |
| Tet18 vs. Tet59 | * | 0.0337 |

**TS4: Statistical summaries for figure 3 -- multiple comparisons testing**

